# Mineralisation of the *Callorhinchus* vertebral column (Holocephali; Chondrichthyes)

**DOI:** 10.1101/2020.07.27.222737

**Authors:** Jacob Pears, Zerina Johanson, Kate Trinajstic, Mason Dean, Catherine Boisvert

**Affiliations:** School of Molecular and Life Sciences, Curtin University, Perth, Australia.; Department of Earth Sciences, Natural History Museum, London, UK.; Max Planck Institute of Colloids and Interfaces, Department of Biomaterials, Am Muehlenberg 1, 14476 Potsdam, Germany.

**Keywords:** Holocephali, *Callorhinchus*, tesserae, mineralisation, evolution, stem group Holocephali

## Abstract

Chondrichthyes (Elasmobranchii and Holocephali) are distinguished by their largely cartilaginous endoskeleton that comprises an uncalcified core overlain by a mineralised layer; in the Elasmobranchii (sharks, skates, rays) this mineralisation takes the form of calcified polygonal tiles known as tesserae. In recent years, these skeletal tissues have been described in ever increasing detail in sharks and rays but those of Holocephali (chimaeroids) have been less well-described, with conflicting accounts as to whether or not tesserae are present. During embryonic ontogeny in holocephalans, cervical vertebrae fuse to form a structure called the synarcual. The synarcual mineralises early and progressively, anteroposteriorly and dorsoventrally, and therefore presents a good skeletal structure in which to observe mineralised tissues in this group. Here we describe the development and mineralisation of the synarcual in an adult and stage 36 elephant shark embryo (*Callorhinchus milii*). Small, discrete, but irregular blocks of cortical mineralisation are present in stage 36, similar to what has been described recently in embryos of other chimaeroid taxa such as *Hydrolagus*, while in *Callorhinchus* adults, the blocks of mineralisation have become more irregular, but remain small. This differs from fossil members of the holocephalan crown group (*Edaphodon*), as well as from stem group holocephalans (e.g., Symmorida, *Helodus*, Iniopterygiformes), where tessellated cartilage is present, with tesserae being notably larger than in *Callorhinchus* and showing similarities to elasmobranch tesserae, for example with respect to polygonal shape.

## Introduction

During ontogeny most vertebrate skeletons are initially composed predominantly of hyaline cartilage and largely replaced by bone via endochondral ossification (Hall, 1975, 2005). In contrast, chondrichthyans, including elasmobranchs (sharks, skates, rays and relatives) and holocephalans (chimaeroids) do not develop osseous skeletons, having secondarily lost the ability to produce endoskeletal bone (Coates et al., 1998; Dean and Summers, 2006; Ryll et al., 2014; Debiais-Thibaud, 2019). Instead, the chondrichthyan endoskeleton remains primarily composed of hyaline-like cartilage, with elasmobranchs developing a comparatively thin outer layer of cortical mineralisation during ontogeny (Hall, 2005; Egerbacher et al., 2006; Dean et al., 2009, 2015; Seidel et al., 2016, 2019; Debiais-Thibaud, 2019). This mineralised tissue begins as small separated islets near the cartilage surface, which gradually grow via mineral accretion to fill the intervening spaces, eventually forming a thin cortex of abutting polygonal tiles called tesserae (Dean and Summers, 2006; Dean et al., 2009, 2015; Seidel et al., 2016, 2019; Dean, 2017). These tiles cover the uncalcified cartilage core and are themselves overlain by a distal fibrous perichondrium (Dean and Summers, 2006; Dean et al., 2009, 2015). This mosaic of uncalcified cartilage, tesserae and perichondrium is called tessellated cartilage and comprises most of the cranial and postcranial skeleton (Kemp and Westrin, 1979; Dean and Summers, 2006; Seidel et al., 2016, 2017a).

Tessellated cartilage is therefore a major component of the skeleton and is currently believed to be a synapomorphy for the entire chondrichthyan group (e.g., Maisey et al. 2019, but see comments therein regarding morphological and histological disparity in stem-chondrichthyans). Contemporary examination of extant chondrichthyan mineralised skeletons and their tissues, however, have almost exclusively focused on sharks (Kemp and Westrin, 1979; Peignoux-Deville et al., 1982; Clement, 1986, 1992; Bordat, 1987, 1988; Egerbacher et al., 2006; Eames et al., 2007; Enault et al., 2016) and rays (Dean et al., 2009, 2015; Claeson, 2011; Seidel et al., 2016, 2017a, b; Criswell et al., 2017a, b). In contrast, mineralised skeletal tissues of extant chimaeroids (Holocephali) have been largely ignored, despite available descriptions of vertebral development and morphology in the late nineteenth to mid-twentieth centuries (Hasse, 1879; Schauinsland, 1903; Dean, 1906); fossil holocephalans have faced similar neglect (e.g., Moy-Thomas, 1936; Patterson, 1965; Maisey, 2013). This has led to contradictory descriptions of chimaeroid tissues (Lund and Grogan, 1997, 2004; Pradel et al., 2009; Dean et al., 2015), prompting calls for more research (Eames et al., 2007; Dean et al., 2015; Enault et al., 2016). Notably, recent examination of chimaeroid mineralised skeletal tissues identified tesseral structures in the vertebral column (synarcual) and Meckel’s cartilage of *Chimaera* and *Hydrolagus* (both Family Chimaeridae; Finarelli and Coates, 2014; Debiais-Thibaud, 2019; Seidel et al., 2019a), seemingly refuting the view that extant chimaeroids lack tessellated cartilage.

In order to address this controversy, and determine whether tessellated cartilage is a shared character among cartilaginous fishes, we examine mineralisation in the skeletal tissue of representatives of a second family of extant holocephalans, the Callorhinchidae, focusing on the synarcual of the elephant shark (*Callorhinchus milii*). The synarcual is a fused element in the anterior vertebral column (Claeson, 2011; Johanson et al., 2013, 2015, 2019; VanBuren and Evans, 2017) and is one of the better anatomical structures for mineralised tissue characterization, being formed early in development and also mineralising early (Johanson et al., 2015, 2019). Synarcual mineralisation progresses from anterior to posterior, and dorsal to ventral, allowing the observation of different mineralisation patterns and stages within a single anatomical structure (Johanson et al., 2015). We report the presence of a layer of mineralisation in the *Callorhinchus* embryo, comprising small, irregularly-shaped units, maintained in adults, and lacking many of the characteristics of tesserae in the elasmobranchs. To provide further phylogenetic context we also examined mineralised tissues in fossil members of the Callorhinchidae (*Edaphodon*; Nelson et al. 2006), as well as stem-group holocephalan taxa (e.g., *Cladoselache*, *Cobelodus*, *Helodus*, Iniopterygiformes; Coates et al., 2017, 2018; Frey et al., 2019). The tesserae in these stem-group holocephalans are larger than in *Callorhinchus*, and more similar in shape to polygonal elasmobranch tesserae. Thus, the evolution of skeletal mineralisation in Chondrichthyes may have involved a progressive reduction of mineralisation in the Holocephali, relative to the elasmobranchs.

## Materials and Methods

### 2.1 Histological sections of *Callorhinchus milii* synarcual

To gain insight into the development of mineralised tissues, slides of the synarcual from a sectioned embryo of an elephant shark (*Callorhinchus millii*; section thickness ~30μm; Life Sciences Department, Natural History Museum, London) were examined by light microscopy using an Olympus BX51 compound microscope and Olympus DP70 camera and management software. These slides were prepared sometime during the 1980s and the animal is estimated to represent stage 36 (near hatching, based on the calculated size of the individual (110–135 mm; Didier et al., 1998). This developmental stage is ideal to study mineralisation as it is small enough to section but mature enough to show mineralisation, including more mineralisation anterodorsally, progressing posteroventrally. This in effect provides ontogenetic information on how mineralisation develops, in one individual.

### 2.2 Adult *Callorhinchus milii*

Two adult female of *C. milii* were captured by rod and reel from Western Port Bay, Victoria, Australia (Permits: RP1000, RP 1003 and RP1112) with the authorisation and direction of the Monash University Animal Ethics Committee (Permit: MAS-ARMI-2010-01) and kept according to established husbandry methods (Boisvert et al., 2015). These specimens died in captivity and were frozen.

### 2.3 Scanning Electron Microscopy

The synarcual of one of these adult *C. milii* specimens was dissected out and either small layers of mineralised tissue or cross sections of the vertebrae were collected. Samples were macerated in a trypsin solution (0.25g Trypsin Sigma T-7409 Type II-S from porcine pancreas in 100mL 10%PBS) and warmed in a 38°C water bath. Samples were extracted from the solution every hour to remove macerated flesh and fascia using scalpels, needles and forceps. This was repeated until sufficient flesh had been removed to observe the mineralised surface. To prevent distortion, samples were placed between Teflon blocks before being air-dried until firm. Cross sections were embedded in a Struers CitoVac using Struers EpoFix Resin and EpoFix Hardener mixed in a 50:6 weight ratio and polished using a Struers Tegramin-30. All samples were given a 3nm conductive coating of pure platinum using a Cressington 208HR sputter coater. Samples were imaged using a TESCAN MIRA3 XMU variable pressure field emission scanning electron microscope (VP-FESEM) using backscatter mode (voltage: 15 kv; working distance: 6–15 mm; Tescan Mira3 VP-FESEM instrumentation, John de Laeter Centre, Curtin University). The remaining adult synarcual (Johanson et al. 2015: fig. 7) was dissected out, air dried and imaged using a FEI Quanta 650 FEG SEM in secondary electron mode (voltage: 10 kv; working distance: 14.7 mm).

### 2.4 Macrophotography

Five fossil holocephalans from the Earth Sciences Department, NHM (NHMUK PV P) were chosen to represent extinct taxa, phylogenetically important with respect to the Callorhinchidae and crown-group holocephalans (Coates et al., 2017, 2018; Frey et al. 2019). These comprised: *Cladoselache* (NHMUK PV P.9285), *Cobelodus* (NHMUK PV P.62281a), *Sibirhynchus* (NHMUK PV P.62316b), *Edaphodon* (NHMUK PV P.10343), and *Helodus* (NHMUK PV P.8212). One specimen preserving mineralised cartilage was chosen from each taxon, and photographed using a Canon EOS 600D camera, EOS Utility. Five to ten images of each specimen were taken at different focal depths and the resultant image stack imported into Helicon Focus (v. 6.8.0) to create images with high depth of focus. These specimens were also photographed using a Zeiss Axio Zoom microscope with camera to provide closeup images; as well, tesseral width was determined using the measurement function in the Zen Pro 2 software accompanying the Axio Zoom microscope (Table 1).

**Table 1:**
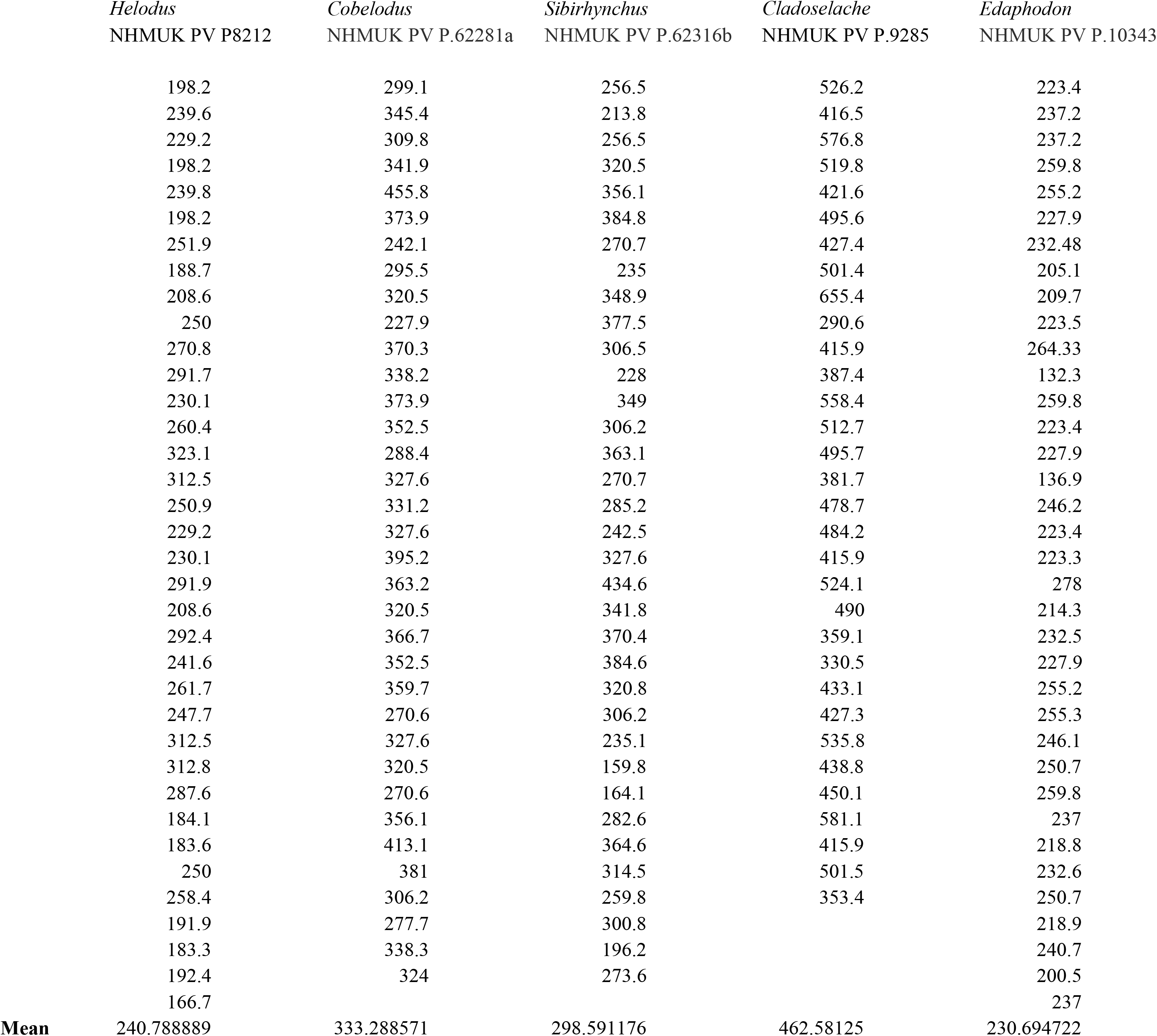
Tesserae width taken from stem group Callorhinchidae *(Edaphodon)* and several stem group Holocephali *(Sibirhynchus, Cobelodus, Helodus, Cladoselache).* Measurements in microns.

The second adult *Callorhinichus milii* synarcual (Johanson et al. 2015) was air-dried and photographed using Zeiss Axio Zoom microscope to illustrate tesseral shape.

## 3 RESULTS

### 3.1 Histology

#### 3.1.1 General Morphology

The axial skeleton of chondrichthyans typically includes a series of dorsal and ventral cartilages, and in the elasmobranchs, centra associated with the notochord (e.g., Dean, 1895, Gadow and Abbott, 1895, Goodrich, 1930; Compagno, 1977; Criswell et al., 2017b). Mineralisation of the axial skeleton takes a variety of forms, recently summarized by Debiais-Thibaud (2019), with the dorsal and ventral cartilages of most species, as well as the outer centrum, composed of tessellated cartilage (Dean and Summers, 2006; Dean et al., 2009; Criswell et al., 2017b; Johanson et al., 2019). Most of the spool-shaped vertebral centrum comprises areolar mineralisation, with substantial variation in patterns of mineralisation between elasmobranch species (Ridewood, 1921; Dean and Summers, 2006; Porter et al., 2007). Holocephalans also possess dorsal and ventral cartilages (e.g., Dean, 1895; Johanson et al., 2012, 2015), but centra do not develop. Instead, the notochord is surrounded by a fibrous chordal sheath, which contains many calcified rings, except in the Callorhinchidae (Patterson, 1965; Didier, 1995). Holocephalans possess a synarcual, absent in elasmobranchs, which is the focus of the following description.

In the *Callorhinchus* embryo examined (stage 36), several tissue layers concentrically surround the notochord. Most proximal is a thin basophilic membrane, the elastic interna, adherent to the outside of the notochord (Figure 1A–D, nc, el.int). Distal to this membrane is a thick (~665μm) fibrous sheath (Figure 1A, B, fb.sh), which is largely composed of spindle shaped cells (Figure 1C, D). Abutting the sheath dorsally and ventrally are separate bilateral pairs of cartilages, the basidorsals and basiventrals, respectively (Figure 1B, D, bv, bd). Immediately dorsal to the sheath is the spinal cavity, containing the spinal cord, which is surrounded ventrolaterally by the basidorsal cartilages and dorsally by the neural arch cartilage (Figure 1B, sp.c, sp.cd, bd, na). Spinal nerves are also visible in section, with the dorsal root exiting the neural tube towards the dorsal root ganglion situated lateral to the vertebral column (Figure 1A, B, d.rt, d.rt.g). The hyaline cartilages associated with the vertebral column—the neural arch, basidorsals and basiventrals— fuse anteriorly to form the synarcual, which surrounds the majority of the fibrous sheath and spinal cavity, while maintaining foramina for the dorsal root (Figure 1A).

**FIGURE 1.**
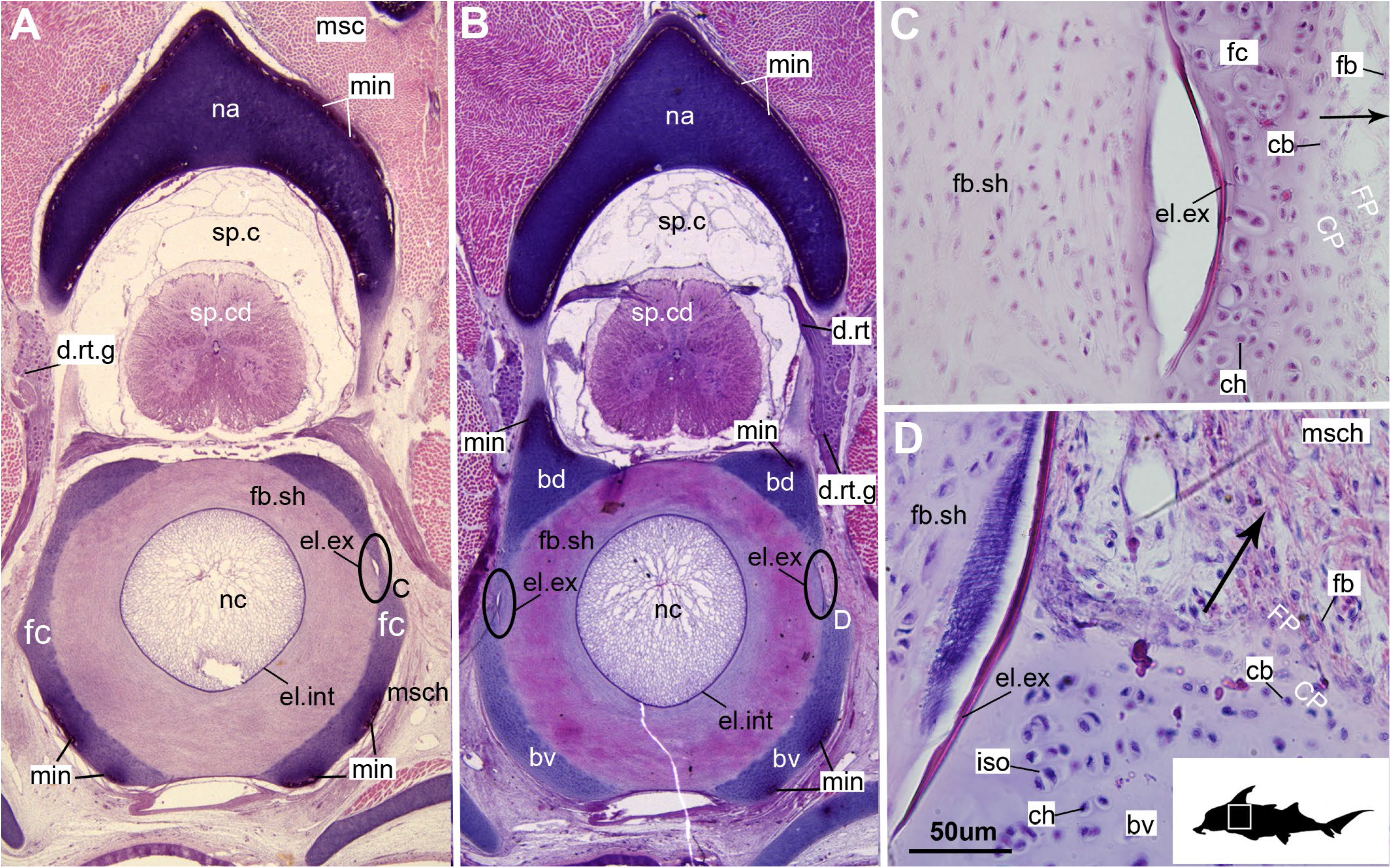
Histological sections through the synarcual (anterior fused vertebrae) of a stage 36 embryo of *Callorhinchus milii* (Holocephali; Callorhinchidae). A, B, section showing neural arch surrounding the spinal cord and basidorsal and basiventral arches surrounding the notochord. C, closeup of region indicated in A; D, closeup of region indicated in B. Black s in C, D indicate direction of appositional growth of the basiventral cartilage. Abbreviations: bd, basidorals, bv, basiventral; cb, chondroblasts; ch, chondrocyte; CP, perichondrium including chondroblast cells; d.rt, dorsal root of the spinal nerve; d.rt.g, dorsal root ganglion of the spinal nerve; el. ex, elastica externa; el.int, elastic interna; fb, fibroblasts; fb.sh, fibrous sheath surrounding the notochord; fc, fused cartilage; FP, perichondrium including fibroblast cells; iso, isogenous group of chondrocytes; min, mineralisation; msc, musculature; msch, mesenchymal cells; na, neural arch, nc, notochord; sp.c, spinal cavity; sp.cd, spinal cord. Black silhouette of *C. millii* indicates approximate region shown in the figure.

In these histological slides, areas of mineralisation are limited to the distal peripheries of the vertebral column-associated cartilages (Figures 1A, B, 2, min). These mineralised tissues are bordered externally by a fibrous perichondrium and a thin, cell-rich layer of cartilage (Figures 2, 3, FP, SC), similar to the supratesseral cartilage in the stingray *Urobatis halleri* (Seidel et al., 2017).

**FIGURE 2.**
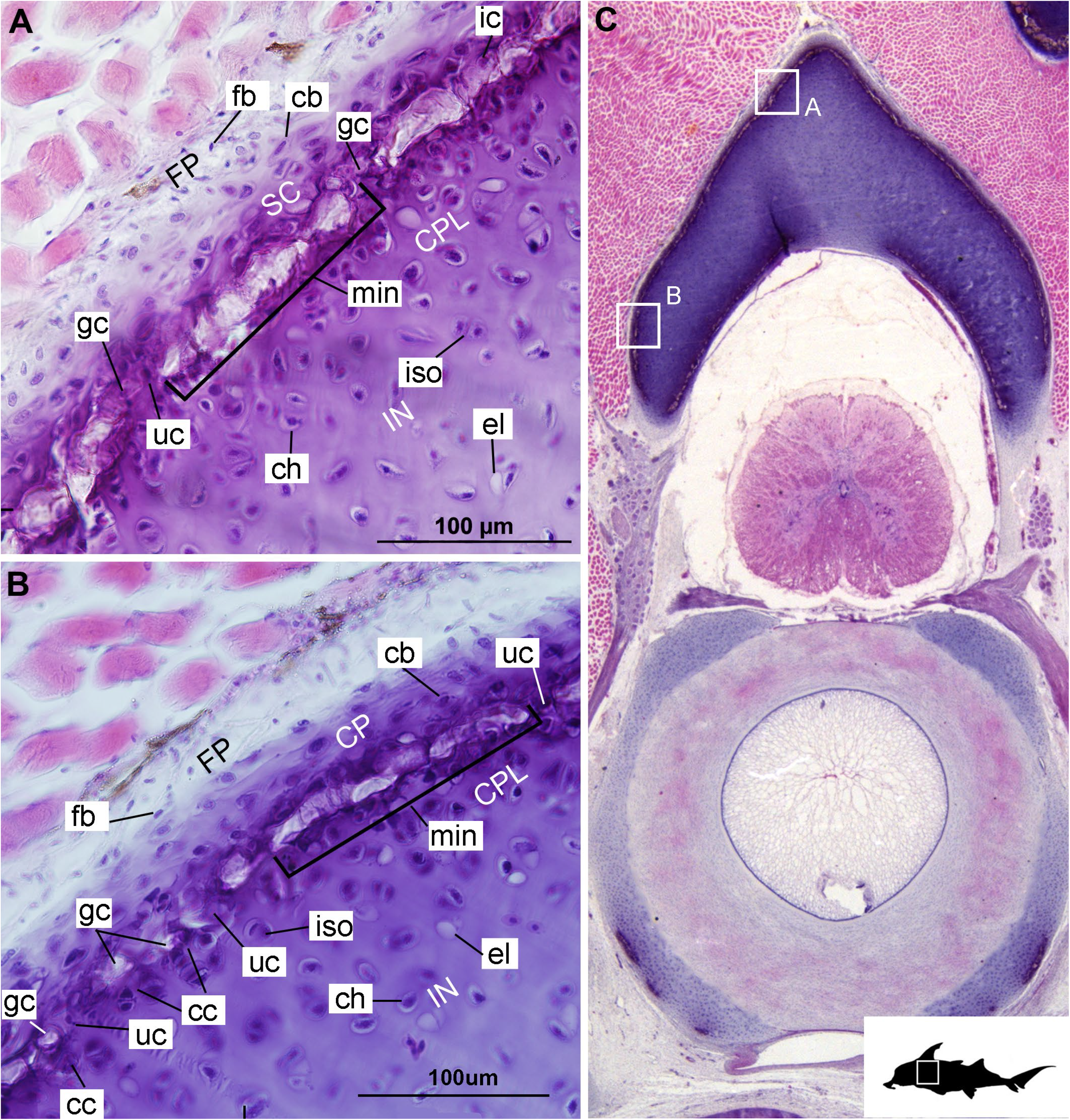
Histological sections through the anterior synarcual (anterior fused vertebrae) of a stage 36 embryo of *Callorhinchus milii* (Holocephali; Callorhinchidae). A, B, closeups showing perichondrium, cartilage and mineralisation in the neural arch; C, overview of section with locations of closeup views indicated by white squares. Abbreviations: As in Figure 1, also cc, clustered chondrocytes CPL, chondrocyte proliferative layer; el, empty chondrocyte lacunae; gc, calcification globule; ic, chondrocyte that is being engulfed or has been incorporated; uc, uncalcified cartilage; IN, internal cartilage; SC, supratesseral/mineral cartilage. Black silhouette of *C. millii* indicates approximate region shown in the figure.

**FIGURE 3.**
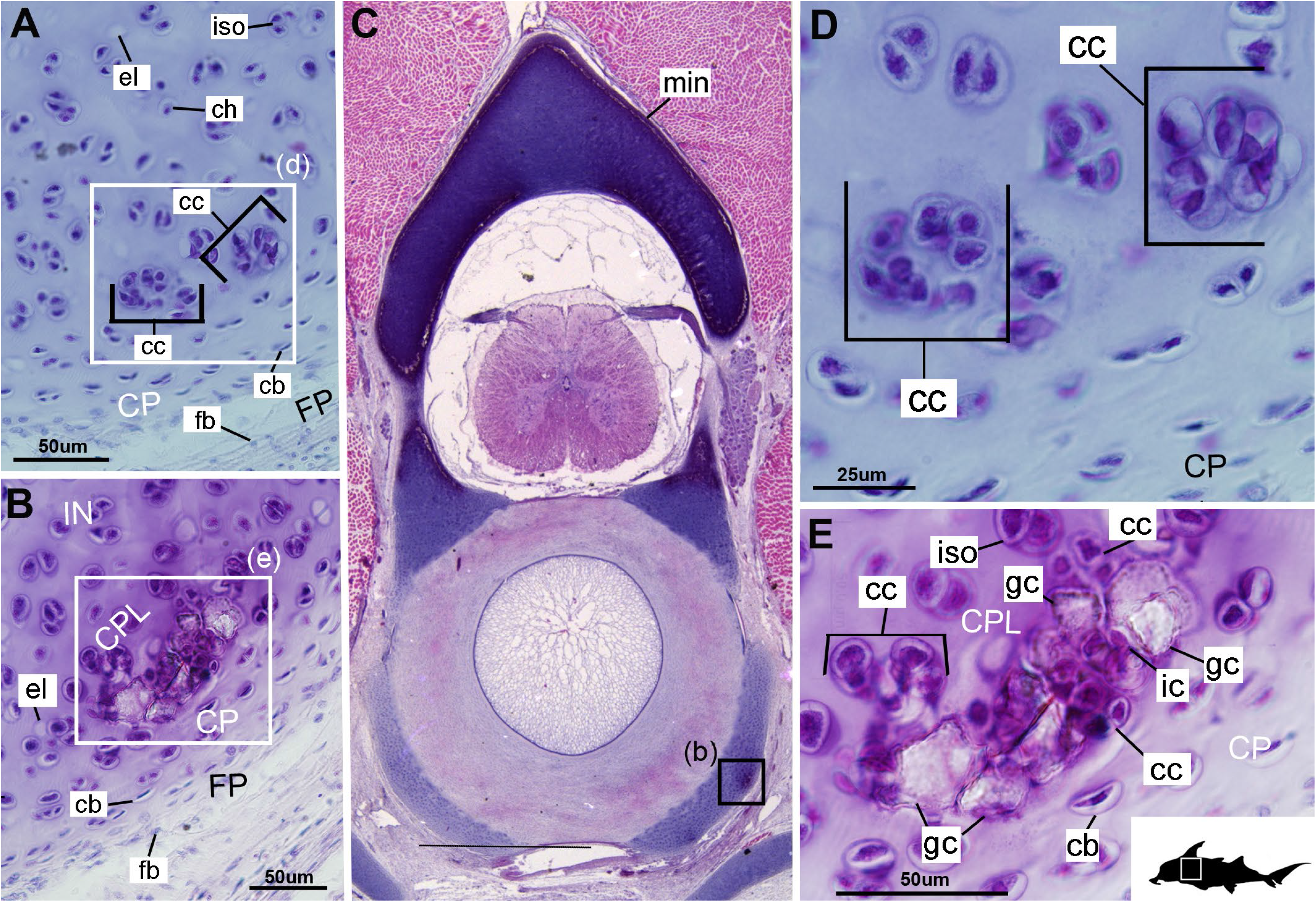
Histological sections through the posterior synarcual (anterior fused vertebrae) of a stage 36 embryo of *Callorhinchus milii* (Holocephali; Callorhinchidae). A, B, closeups showing initial mineralisation in a basiventral and clustered chondrocytes in the same location in the preceding section; C, overview with location of closeup view of initial mineralisation indicated by a black square; D, E, close ups of A, B, locations indicated by white squares. Abbreviations: As in previous Figures, also dm, developing mineralisation. Black silhouette of *C. millii* indicates approximate region shown in the figure.

#### 3.1.2 Cellular Aspects

Cells within the neural arch, basidorsal and basiventral cartilages can be categorised with respect to morphology, and distribution within the cartilage. Chondrocytes located deeper in the cartilage interior (Figures 2A, 3B, IN, ch) are approximately similar in terms of cell morphology and density: ≥200μm from the periphery, cells are sparsely distributed within the intracellular matrix, with most being ovoid in shape and located in open circular spaces identified as lacunae (diameter: ~15 μm). Chondrocytes often occur in pairs, which may indicate recent mitotic activity (Figures 2A, B, 3A, ch; Kheir and Shaw, 2009). This deeper (≥200μm from the periphery), interior cartilage also contains relatively greater quantities of empty lacunae compared to the more peripheral cartilage (Figures 2A, B, 3A, el), which may indicate chondroptosis (chondrocyte apoptosis; Roach et al., 2004), although this is normally associated with chondrocyte hypertrophy, which has not been observed in chondrichthyans (Dean et al., 2015; Seidel et al., 2017b; but see Debiais-Thibaud [2019] for a summary of contrary opinions). Closer to the periphery, within ≥100 μm of the outer edge, and immediately proximal to mineralised tissue, chondrocytes are clustered within a distinct layer (Figures 2, 3B, CPL) and appear uniformly ovoid. This area displays a greater variation in cell size as it contains many smaller chondrocytes (diameter: 5–10 μm), and fewer empty lacunae compared to the interior. In addition, this area contains notably more isogenous groups relative to the interior, which may indicate higher rates of chondrocyte proliferation (Figures 2A, B, 3B, E, iso; Kheir and Shaw, 2009). In some regions, mineralisation is absent at the periphery (i.e., there is no tesseral layer); in these areas, cell distribution and morphology are more similar to the interior (Figure 3A).

There appears to be a gradient of cell morphology from the perichondrium, through the supratesseral cartilage, potentially relating to a cellular transition from fibroblasts cells in the perichondrium, differentiating to chondroblasts in the supratesseral cartilage, and then to chondrocytes in the main body of the cartilage as described in Genten et al. (2009; Figures 1C, D, 2A, B, CP, FP, fb, cb, ch). These cartilages seem to display interstitial and appositional growth, with the former involving the mitotic division of single chondrocytes into a cluster of cells (the isogenous groups), and matrix deposition between these to increase the size of the element (e.g., Figures 1D, 2A, B, iso), while the latter occurs at the cartilage margin through chondrocyte differentiation and matrix deposition (Figure 1B–D; Hall, 2005; Kheir and Shaw, 2009). For example, in several sections, the basidorsal and basiventral cartilages are still separate (e.g., Figure 1B), but appear to show a region of appositional and interstitial growth or matrix deposition at their margins (Figure 1B–D, black arrows showing direction of growth).

#### 3.1.3 Mineralisation

In the neural arches, the distribution of mineralised tissue is more complete, extending along almost the entire periphery excluding only the ventro-mesial concave part of the arch (Figures 2C, 4B). Within the basiventrals, mineralised tissue is also found near the periphery, but by comparison to the neural arches, is only patchily distributed (Figures 3, 4), with individual units more variable and irregular in shape (Figures 3B, E, 4, min). In the neural arches, these units are more rectangular and flatter (Figures 2A, B, 5A, B). Nevertheless, mineralised tissues in all vertebral elements lack a regular geometry and any differentiation into inner and outer regions. Additionally, beyond being limited to the cartilage periphery beneath the fibrous perichondrium, these tissues lack any organisation, reflecting the lack of a regular geometric shape to the individual units.

**FIGURE 4.**
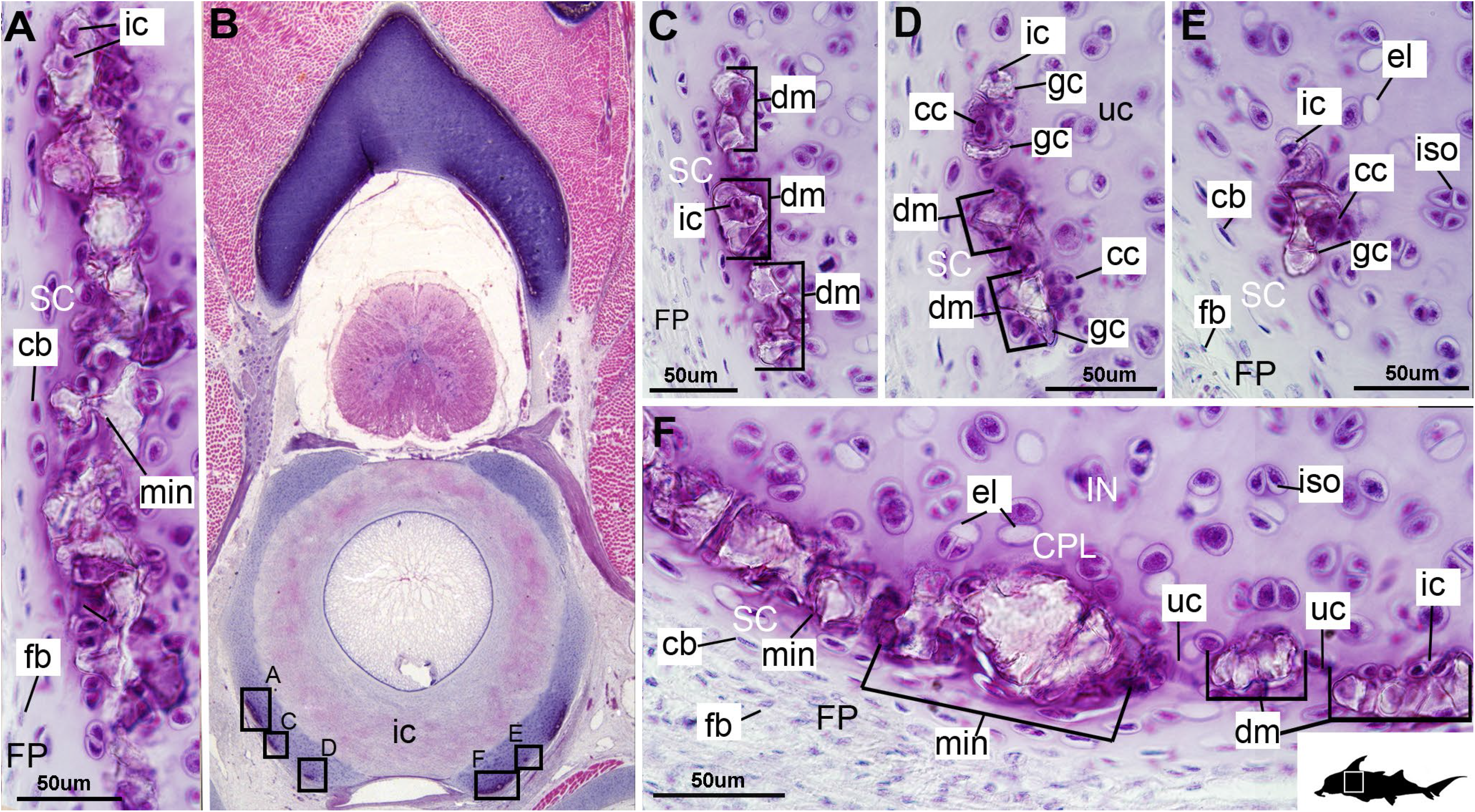
Histological section through the anterior synarcual (anterior fused vertebrae) of a stage 36 embryo of *Callorhinchus milii* (Holocephali; Callorhinchidae). A, C, D, E, F, closeups showing perichondrium, cartilage and mineralisation in basiventrals; B, overview of section through the neural arch with locations of closeup views indicated by black squares. Abbreviations: As in previous Figures. Black silhouette of *C. millii* indicates approximate region shown in the figure.

**FIGURE 5.**
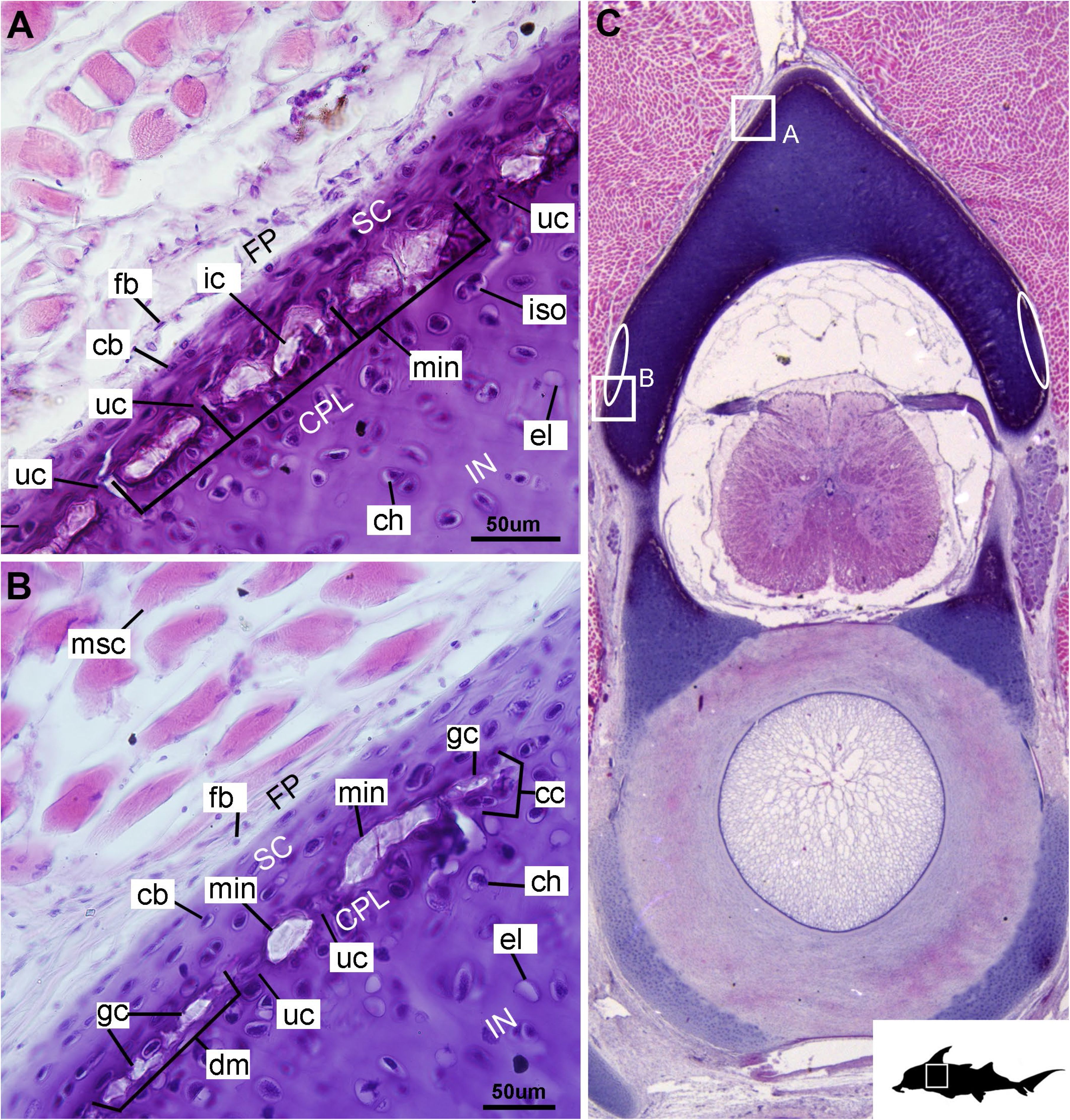
Histological section through the posterior synarcual (anterior fused vertebrae) of a stage 36 embryo of *Callorhinchus milii* (Holocephali; Callorhinchidae). A, B, closeups showing perichondrium, cartilage and mineralisation in the neural arch; C, overview of section with locations of closeup views indicated by white squares. Abbreviations: As in previous Figures. Black silhouette of *C. millii* indicates approximate region shown in the figure.

Tissue mineralisation appears to be preceded by the clustering of chondrocytes within cartilage below the perichondrium (Figures 3A, D, 5B, cc, FP), because amongst these clustered chondrocytes, the inception of mineralisation can be observed via the formation of small islands of calcification (≤25 μm). Initially, these partially encircle these cells (Figures 3B, E, 5B, gc, ic), but come to surround the chondrocytes, forming a thin (≤50 μm) layer of irregular units of mineralised tissue (Figures 4C, D, 5A, B, dm, min, ic). The anteroposterior order of this sequence suggests that the initial islands expand, through the calcification of the surrounding tissue. Alternatively, these islands may grow by fusing together to form larger units, as indicated by the presence of small mineralisation foci between units (Figure 6). More anteriorly, the irregular units (~50–150 μm wide, in cross section, variable in shape), are acellular, suggesting the mineralisation eventually completely engulfs the chondrocytes, resulting in their death. We identify these units as tesserae, with comparison of these mineralised units to the tesserae of sharks and rays discussed further below.

**FIGURE 6.**
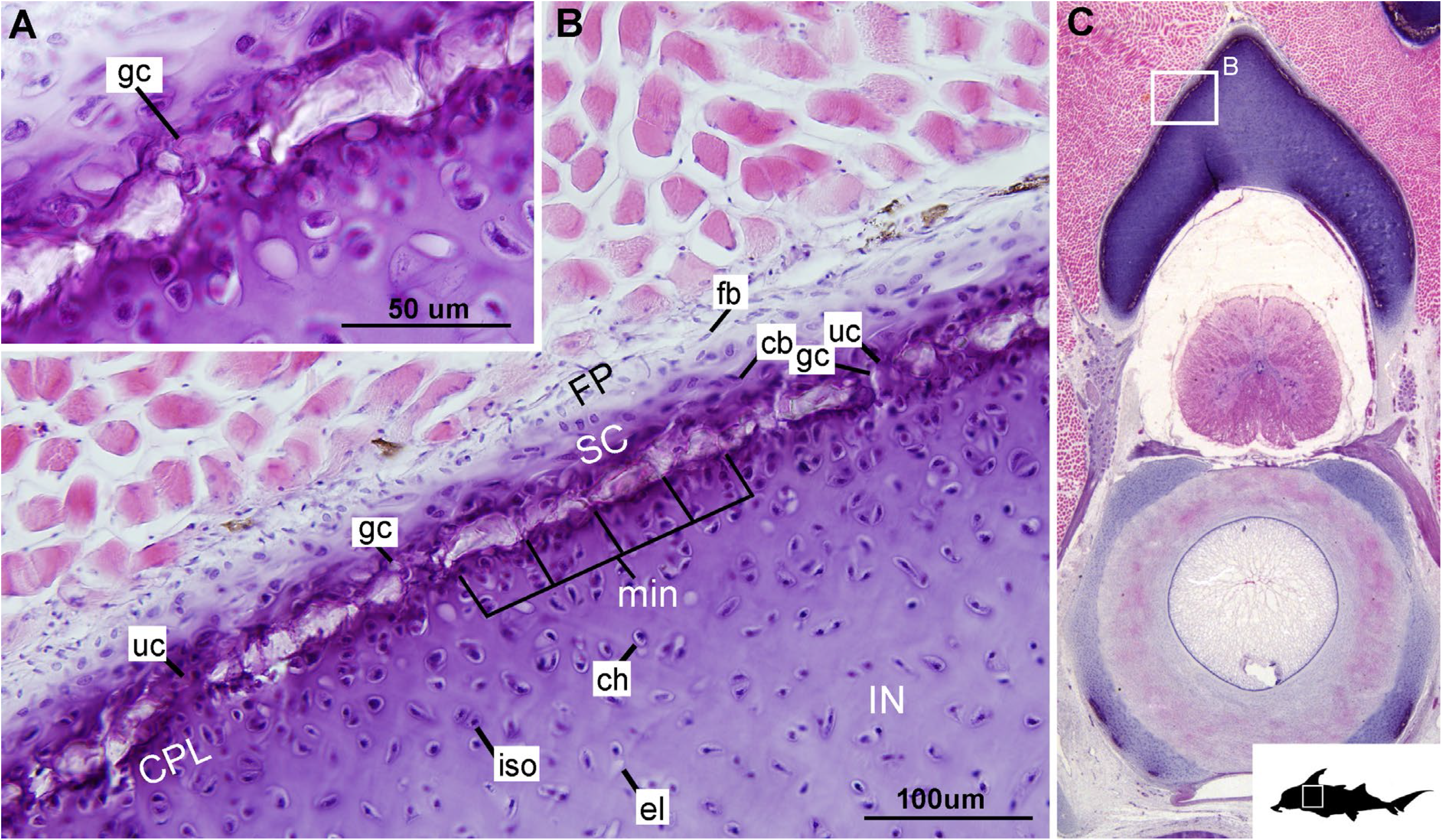
Histological section through the synarcual (anterior fused vertebrae) of a stage 36 embryo of *Callorhinchus milii* (Holocephali; Callorhinchidae). A, B, closeups showing the potential formation of new mineralisation foci between already existing units; C, overview of section through the neural arch with locations of closeup views indicated by white squares. Abbreviations: as in previous Figures. Black silhouette of *C. millii* indicates approximate region shown in the figure.

The presence of less mineralised tissue in posterior sections (Figures 3, 5), compared to anterior (Figures 2, 4), indicates that mineralisation progresses anteroposteriorly along the vertebral column. Thus, tesserae are smaller in more posterior sections (50–100 μm) and more regularly separated by regions of unmineralised cartilage (Figures 4F, 5A, B, min, uc), reflecting their earlier developmental stage.

### 3.2 Scanning Electron Mircoscopy (SEM)

In planar views of the external surface of the synarcual of adult *Callorhinchus milii*, the mineralised layer appears to comprise a tessellated surface of irregular tiles that are separated by (~5um) thin strips of uncalcified cartilage (Figure 7A, B min, uc). These tesserae do not have a uniform shape or size, ranging from 50–150μm in width (Figures 7B; Supplementary Info Figure 1). From this perspective, these tesserae do not appear to possess features that are currently considered common among elasmobranch tesserae (e.g., mineralised ‘spokes’ at the contact points between tesserae, intertesseral joints, vital chondrocytes; reviewed in Seidel et al., 2016, 2019b).

**FIGURE 7.**
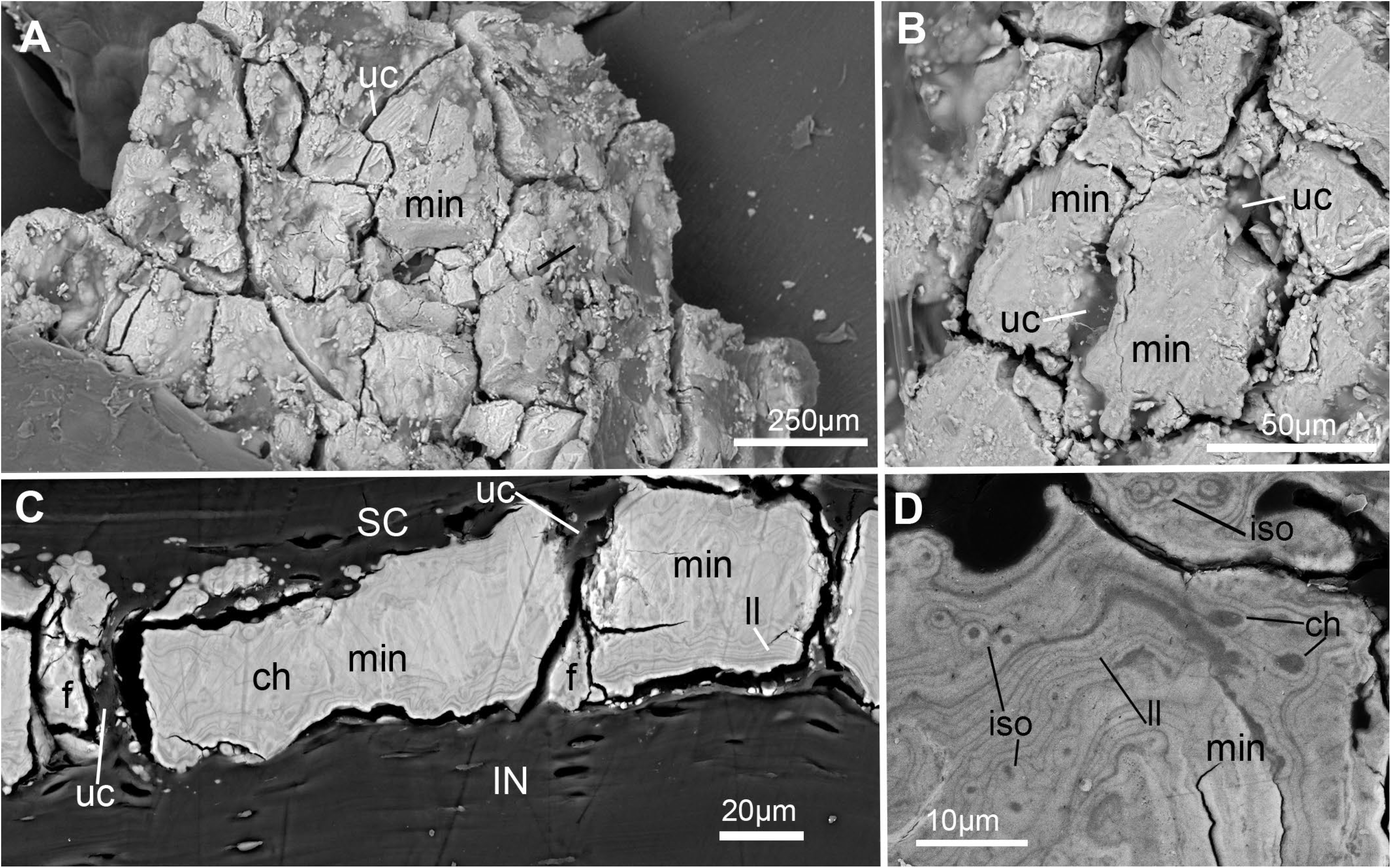
SEM images of mineralisation from the synarcual (anterior fused vertebrae) of an adult *Callorhinchus milii* (Holocephali; Callorhinchidae). A, overview of tesselated mineralisation from a planar perspective; B, close up of mineralisation from a planar perspective; C, mineralisation from a transverse perspective; D, close up of mineralisation surface in transverse perspective; Note brightness and contrast of C, D have been altered to more clearly visualise morphology. Abbreviations: As in previous Figures, also f, fragments ll, Liesegang lines.

In transverse view, the mineralised tissue forms a single layer of tesserae, tightly arranged units of irregular blocks separated by very thin (<5 μm) strips of uncalcified cartilage (Figure 7C, min, SC, IN, uc). In this perspective, the tesserae are 30–50 μm thick and 30–150 μm wide (Figure 7C). Some cracking of the mineralisation during sample preparation is visible, but individual tesserae can be identified by comparing and matching Liesegang lines between adjacent fragments (Figure 7C, ll, f). Liesegang lines are concentric, wave-like patterns of varying mineral density visible in the mineralised tissue, and are particularly prominent near the lateral margins of the mineralised units (Figure 7C, D, ll).

Spheroidal mineralised regions, surrounded by Liesegang lines and approximately the size and shape of chondrocytes also permeate the tesserae (Figure 7D, ch). These are likely calcified (micropetrotic) cells, are variously sized (~1–5 μm), and appear to be organised in clusters (isogenous groups), suggesting some have been calcified during mitosis (Figure 7D, ch, iso).

### 3.3 Mineralisation in stem Holocephali and fossil Callorhinchidae

Following phylogenetic review Coates et al. (2017, 2018; Dearden et al., 2019; Frey et al., 2019), several taxa that were previously resolved as stem group chondrichthyans (basal to the clade Elasmobranchii + Holocephali; Pradel et al., 2011), are now resolved as stem holocephalans, joining more crownward stem holocephalans including the Iniopterygiformes (Zangerl and Case, 1973), *Helodus* (Moy-Thomas, 1936), *Kawichthys*, *Debeerius* and *Chondrenchelys*, the latter being the sister taxon to the crown group Holocephali (chimaeroids) (Figure 8). Tessellated calcified cartilage has been variously identified among these stem-group Holocephali: this includes taxa assigned to the Symmorida, such as *Dwykaselachus* (Coates et al., 2017: extended data figure 1d), *Ozarcus* (Pradel et al., 2014), *Cladoselache* (“minute granular calcifications”; Dean, 1894; Figure 9A), *Akmonistion* (“prismatic calcified cartilage” Coates and Sequiera, 2001), *Damocles* and *Falcatus* (Lund and Grogan, 1997), also present in *Cobelodus* (Figure 9B). In all of these taxa, the tesselated layer is comprised of recognizable polygonal units, although in *Cladoselache*, the edges of the units appear less regular. This may represent the presence of mineralised ‘spokes’ extending between the tesserae: spokes are hypermineralised tissue regions associated with points of contact between elasmobranch tesserae, often represented externally by lobulated extensions along tesseral margins (Seidel et al., 2016; Jayasankar et al. 2020). Such structural extensions, suggestive of mineralised spokes, are even more clearly present in *Cobelodus* (Figure 9C).

**FIGURE 8.**
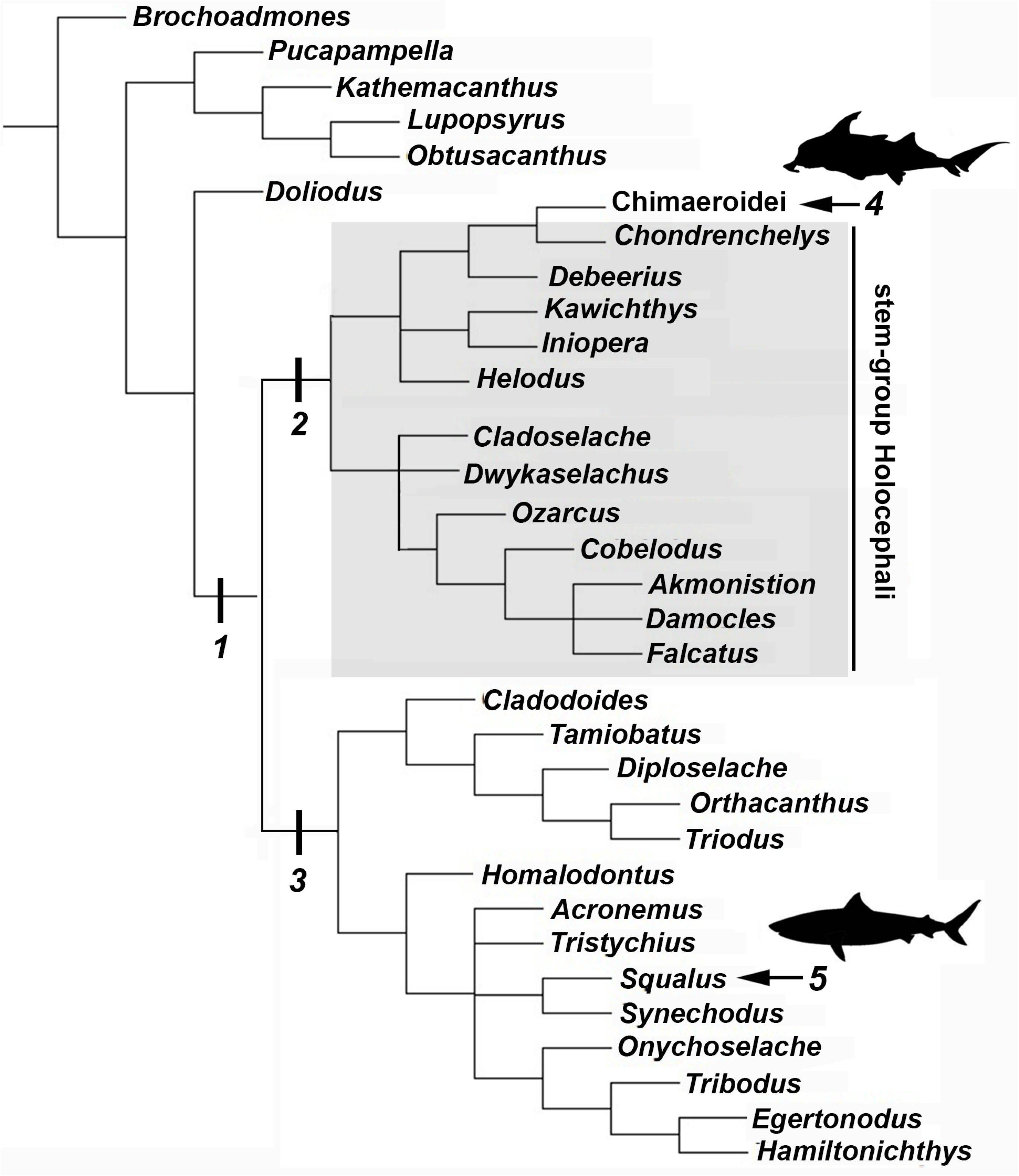
Chondricthyan phylogeny (after Coates et al. 2018). 1, Crown–group Chondrichthyes (Holocephali + Elasmobranchii); 2, Holocephali; 3, Elasmobranchii; 4, Crown–group Holocephali; 5, Crown–group Elasmobranchii.

**FIGURE 9.**
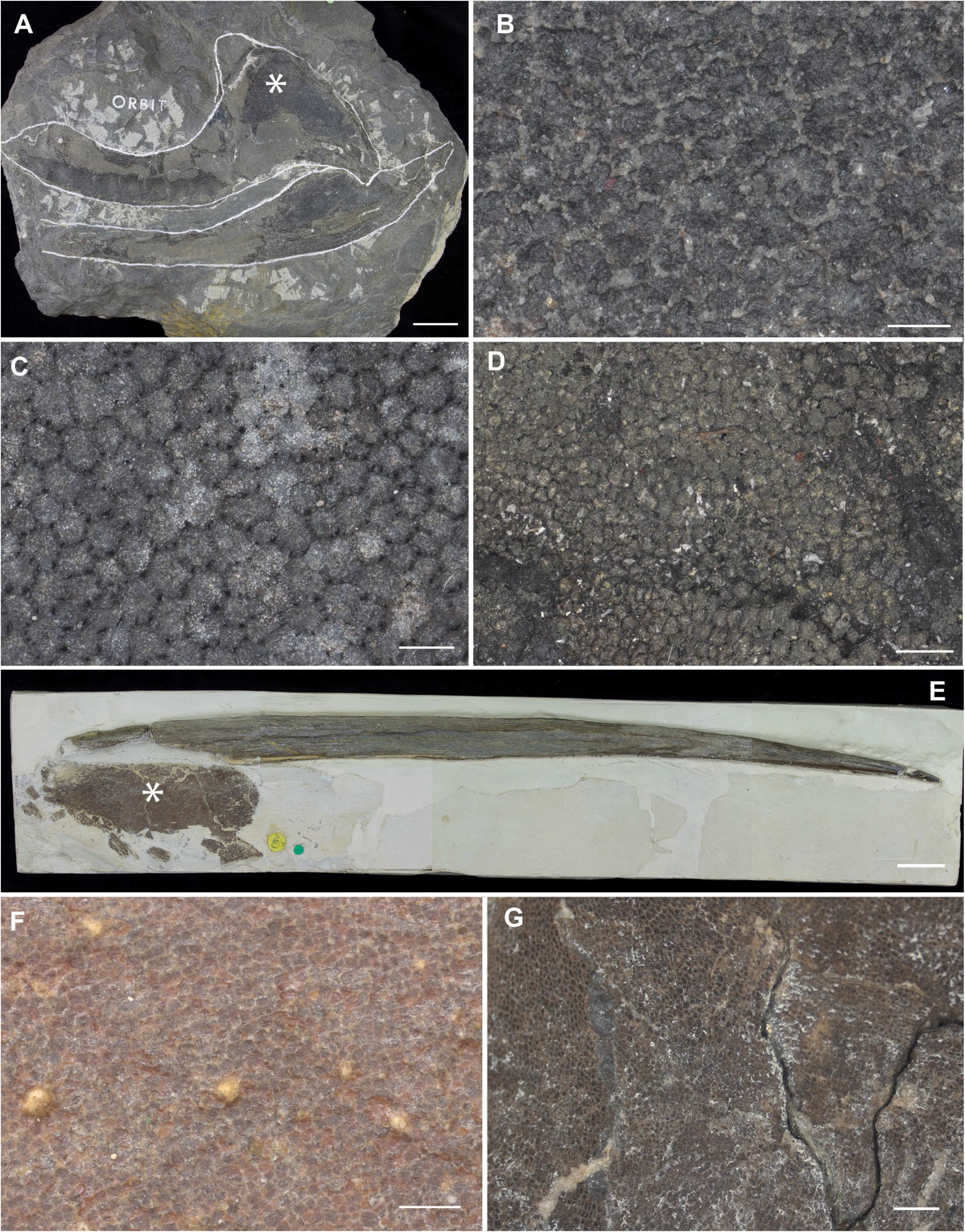
Mineralised cartilage in stem Holocephali and crown group Holocephali (Fig. 8). A, NHMUK PV P.9285, *Cladoselache*, stem Holocephali, palatoquadrate and Meckel’s cartilage; asterisk indicates area shown in B; B, polygonal mineralisation (tesserae), with irregular margins; C, NHMUK PV P.62281a, *Cobelodus*, stem Holocephali, more regular polygonal mineralisation (tesserae); D, NHMUK PV P.62316b, *Sibirhynchus*, stem Holocephali, polygonal mineralisation (tesserae); E, F, NHMUK PV P.10343, *Edaphodon*, Family Callorhinchidae, crown group Holocephali (Fig. 8, 2), E, dorsal fin endoskeletal support (with dorsal fin spine, anterior to left), asterisk indicates area shown in F; F, closeup showing polygonal mineralisation (tesserae); G, NHMUK PV P.8212, *Helodus*, polygonal mineralisation. See Table 1 for tessera sizes in these taxa.

In the more crownward stem holocephalans, comparable polygonal tesserae are also present, including in *Kawichthys* (“tesserate prismatic calcified cartilage”, Pradel et al., 2011) and the Iniopterygiformes (“calcified cartilage prisms”, Zangerl and Case, 1973), represented by *Sibirhynchus* in Figure 9D. Particularly small tesserae (Table 1) are present in *Helodus* (“minute tesserae”, Moy-Thomas, 1936; Figure 9G), and *Chondrenchelys* (“tessellated calcified cartilage”, Finarelli and Coates, 2014: fig. 7B). There appears to be more variation in the shape of these polygons, and signs of mineralised spokes are less apparent in these taxa, but this may be due to postmortem distortion. With respect to the fossil taxa assigned to the Callorhinchidae (crown group Holocephali), mineralised tissue units in *Edaphodon* appear to maintain a polygonal shape, compared to the stem holocephalans just described (Figure 9E, F). The width of tesserae in these fossil taxa was measured (Table 1) for comparison to the size of mineralised units in adult *Callorhinchus*; (50–150μm, as noted above) the tesserae of all fossil taxa were notably larger than in *Callorhinchus*, discussed further below.

## 4 Discussion

### 4.1 Mineralised Tissue Development

Currently, the ultrastructure and ontogeny of mineralised endoskeletal tissues of chimaeroids is poorly described, with previous work only providing a broad overview of developmental trajectories, showing that mineralisation in the vertebral skeleton (in the synarcual) progresses from anterior to posterior and dorsal to ventral, as indicated by a micro-CT scan of a *Callorhinchus milii* adult (Johanson et al., 2015; Figure 10A). The series described above in the stage 36 embryo goes beyond this to capture fine histological detail related to the progression of this mineralisation.

**FIGURE 10.**
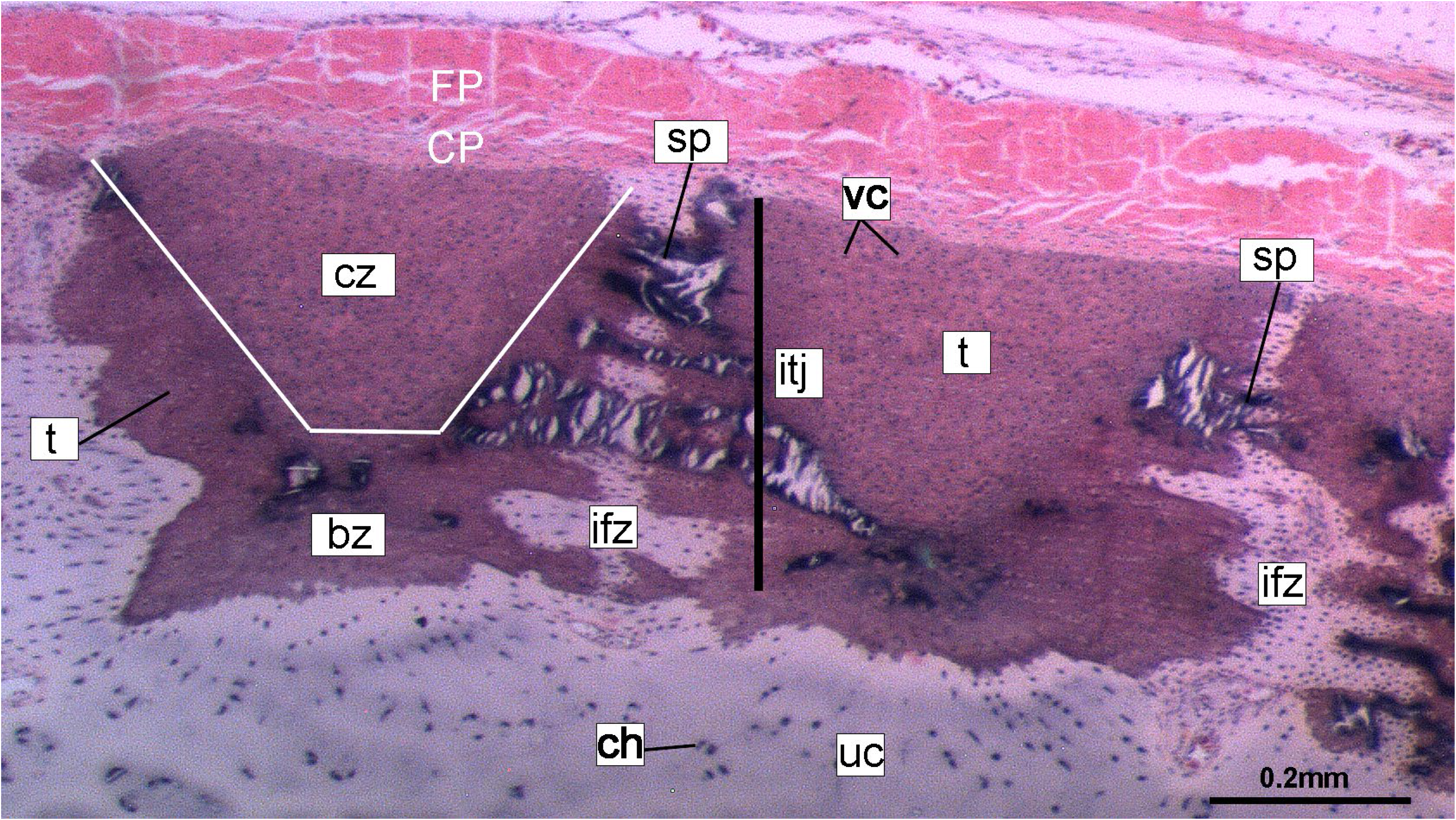
Histological section of tessellated cartilage of a batoid ray (*Raja*). Abbreviations: as in previous Figures also bz, body zone; CP, chondrogenic perichondrium cz, cap zone; itj, intertesseral joint; FP, fibrous perichondrium; sp, spoke; t, tesserae; vc, vital chondrocytes

The development of mineralised tissue described here for *C. milii* shares some similarities with the development of elasmobranch tesserae (Seidel et al., 2016; Debiais-Thibaud, 2019). Tesserae in elasmobranchs such as the batoid ray *Urobatis halleri* initially develop as patches of globular mineralisation interposing within clusters of flattened, subperichondral chondrocytes (at a distance from the perichondrium). These chondrocytes become entombed by the growth of these mineralised intra-chondrocyte septa, by mineral accretion (Dean et al., 2009; Seidel et al., 2016). This accretion and entombment process is similar to the inception of mineralisation observed in *C. milii* and elasmobranch mineralised septae bear resemblance in shape and size to the developing mineralisation of *C. milii* (Figures 4C, D, 5A, B). Through development of *U. halleri*, these mineralised septae continue to grow and engulf chondrocytes, eventually forming discrete, but abutting tesserae, which contain vital chondrocytes and closely border the perichondrium (Seidel et al., 2016; also in the batoid *Raja clavata*, Debiais-Thibaud, 2019). In a general sense, elasmobranch tesserae are not dissimilar to some of the developing units of mineralisation in *C. milii* (Figure 4F, dm, ic), which also border the perichondrium, grow via calcification of surrounding cartilage matrix and, at least early in development, contain chondrocytes which appear to be vital. Additionally, mineralised tissues in *C. milii* appear to be overlain by a distinct layer of uncalcified cartilage, beneath the perichondrium (Figure 2A, SC). This resembles the thin layer of ‘supratesseral uncalcified cartilage’ intervening between tesserae and perichondrium in elasmobranchs such as *U. halleri* and *Scyliorhinus canicula* (Bordat, 1988; Egerbacher et al., 2006; Enault et al., 2015; Seidel et al., 2016, 2017a, Debiais-Thibaud, 2019, contra Kemp and Westrin, 1979).

Despite these similarities, mineralisation in adult *C. milii* appears to be a distinct form of tessellated calcified cartilage. From a planar perspective, the mineralised tissue seems to comprise a more irregular mosaic of tesserae than typically seen in elasmobranchs, with near-abutting calcified tiles separated by uncalcified cartilage (Figure 7A, B, min, uc; Supplementary Info Figure 1). From a transverse perspective, these tissues are similar to that of embryo, arranged as a single layer of tightly arranged units separated by very thin strips of uncalcified cartilage and sandwiched between a supratesseral layer and internal uncalcified cartilage (Figure 7C, D, min, uc, SC, IN). From this perspective, the tesserae of *C. milii* most closely resemble the tesserae of the sevengill shark (*Notorynchus cepedianus*) in terms of their arrangement, being very tightly organised and separated by minimal uncalcified cartilage, while also lacking vital chondrocytes; however, they are not comparable in size, being ~19–57% of the size (Seidel et al., 2016). This is notable, as the Hexanchiformes, to which *N. cepedianus* belongs, are considered one of the most primitive of modern selachian groups (Barnett et al., 2012; Tanaka et al., 2013; da Cunha et al., 2017). However, despite this resemblance, it is likely that this tissue organisation does not represent a plesiomorphic trait given the morphology of the skeletal tissues of stem holocephalans such as *Cobelodus* (see section 4.2).

Despite similarities in the early stages of development between elasmobranch and *C. milii* tesserae (e.g., with early growth surrounding vital chondrocytes; Figures 3B, 4E, 5B), the tesserae of *C. milii* differ significantly from those elasmobranchs, and particularly batoids, in terms of size and ultrastructure. The tesserae observed in the stage 36 embryo and adult *C. milii* (Figures 2, 4, Supplementary Info Figure 1, min) are much smaller compared to most elasmobranch tesserae that have been examined (Seidel et al., 2016; Figure 10, t), generally being less than 50 μm thick and ranging from 50–150 μm in width (Figures 2A, 7C), comparable in size to the ~100 μm tesserae of the catshark *Scyliorhinus* (Egerbacher et al., 2006; Seidel et al, 2016; Debiais-Thibaud, 2019). Additionally, *C. milii* tesserae display no internal regionalisation into the cap and body zones (regions in elasmobranch tesserae delineated by cell shape and collagen type, Figure 10, bz, cz; Kemp & Westrin, 1979; Seidel et al, 2016; Chaumel et al, 2020), with no differences observed between the surfaces closer to the fibrous perichondrium, and surfaces surrounded by hyaline cartilage (e.g., Figures 6A, 7C). Chimaeroid tesserae also apparently lack the intertesseral joints and mineralised spokes characteristic of elasmobranch tesserae (Figure 10, itj, sp; Seidel et al, 2016), as well as the Sharpey’s fibres that extend from the perichondrium into the tesserae cap zone in elasmobranchs (e.g. Kemp and Westrin, 1979; Peignoux-Deville et al., 1982; Clement, 1992; Summers, 2000; Seidel et al., 2017). With respect to growth, the presence of Liesegang lines parallel to tesseral edges (Figure 7C, D, ll) suggests calcification in *C. milii* accretes at the margins of tesserae (Figures 2–5, min, dm, gc) in the same manner as elasmobranch tesserae. Additionally and/or alternatively, *C. milii* tesserae may grow through the development and fusion of new, smaller mineralisation foci between existing tesserae (e.g., Figure 6A). Indeed, the irregular and less concentric arrangement of Liesegang lines in *C. milii* tesserae relative to those in elasmobranchs may be indicative of a more multimodal and/or haphazard form of growth, perhaps explaining the varied shape of the observed tesserae (Figure 7).

As noted, chondrocytes appear to be engulfed during mineralisation in *C. milii* (Figures 3B, E, 4E) and may be vital in early stages (Figures 4C, D, F, 5, dm, ic). However, more developed tesserae in embryos and adults appear to be acellular (Figures 2, 4A, 6, min) as any previously entombed chondrocytes appear to have calcified (Figure 7D, cc), a major difference when compared to most elasmobranch tesserae (Seidel et al., 2017; Debiais-Thibaud, 2019; Figure 10). The absence of vital chondrocytes in the tesserae of *C. milii* may have important implications for their maintenance. In batoids, chondrocytes entombed in tesserae (Figure 10) remain vital in uncalcified lacunar spaces and form passages not unlike the canalaculi found in bone (Dean et al., 2010; Seidel et al., 2016; Chaumel et al., 2020). These chondrocytes and the networks they form are thought to have important functions with regard to maintaining the endoskeleton by communicating information about the mechanical environment in a manner similar to osteocytes in bone (Dean et al., 2010; Seidel et al., 2016; Chaumel et al., 2020). Thus, vital chondrocytes are absent in the tissues of the adult and the anterior older (anterior) regions of the synarcual, suggesting these are lost during ontogeny, along with their associated putative mechanosensory networks, and that these functions are either absent or achieved through alternative means.

### 4.2 Chimaeroid Endoskeleton: Form Across Phylogeny

In contrast with elasmobranchs, the mineralised components of the chimaeroid endoskeleton have been subjected to little study. The limited literature available proposes conflicting forms of endoskeletal mineralisation: tesselated calcified cartilage akin to that of elasmobranchs (Hasse, 1879; Seidel et al., 2019b), smooth superficial sheets of continuous calcified cartilage formed from the fusion of tesserae during ontogeny (Lund and Grogan, 1997; Grogan and Lund, 2004; Pradel et al., 2009; Grogan et al., 2015), or a granular texture (*Hydrolagus*, Finarelli and Coates, 2014). Recent histological data from the synarcual of a sub-adult (20 cm) *Hydrolagus* illustrates two forms of mineralised tissue (Debiais-Thibaud, 2019: 116). This includes small (≤50 μm) subperichondral tissues “reminiscent of globular mineralisation” at the periphery of the vertebral body and neural arch that appear to follow a tessellated pattern, and a more irregular form of globular mineralisation deep within the vertebral body surrounding the fibrous chordal sheath.

Based on these few recent reported data on chimaeroid mineralisation, tesserae in *C. milii* seem to share similarities with those of *Hydrolagus*. In both taxa, mineralisation is tessellated and limited to the periphery of structures composed of hyaline cartilage, including the neural arches, basidorsals and basiventrals (vertebral body), though *C. milii* lacks the second deeper layer of globular mineralisation (Debiais-Thibaud, 2019: fig. 6.1). The mineralised tissues of *Hydrolagus* also take the form of small, irregular acellular units, lacking clear separation into upper cap and lower body zones (Seidel et al., 2016; Debiais-Thibaud, 2019). Likewise, in *Chimaera*, mineralisation more clearly takes the form of tesserae, although differences with respect to the more developed batoid tesserae have been described (Seidel et al., 2016).

These few recent descriptions of mineralisation in modern chimaeroids, including that provided here for *C. millii*, indicate that these taxa do not possess sheets of continuous calcified cartilage, nor a granular texture (Lund and Grogan, 1997; Grogan and Lund, 2004; Pradel et al., 2009; Finarelli and Coates, 2014). Instead they appear to support more historical claims (Hasse, 1879) that these organisms possess tesselated skeletal tissues, though contrary to these sources, these are distinctly different from most elasmobranch tesserae. These discrepant accounts may arise from the tissue arrangements; in taxa such as *C. milii* the tesserae are very tightly arranged, being separated by very thin portions of uncalcified cartilage, which may give the impression that the surface comprises of a sheet. The tesserae themselves are covered in a type of fascia (see Materials and Methods, above), which could account for the observations of a granular texture.

By comparison, in a series of stem holocephalans, cartilage mineralisation in what are presumed to be adults occurs as small polygonal units that are very similar among disparate taxa (Figure 9). The polygonal shape is more comparable to tesserae in the Elasmobranchii, including the suggested presence of mineralised spokes at tesseral joints in taxa such as *Cobelodus*. In contrast, polygonal tesserae are more irregular in shape, and spokes appear absent, in more crownward taxa, including *Edaphodon* (Callorhinchidae), a member of the crown group Holocephali. In these features, the tissues of crownward taxa bear the closest resemblance to those of *C. milii* (Callorhinchidae), however, tesserae in *C. milii* are more irregular in shape and much smaller than the irregular polygonal tesserae of *Edaphodon* (Table 1). The presence of shape and structural features in these fossil taxa that echo those in modern elasmobranch tesserae suggests that substantial changes have occurred in mineralisation in living chimaeroids, with a loss of many characteristics of tesserae in other chondrichthyans.

## 5.0 Conclusion

Whilst tessellated cartilage has been suggested to be a shared characteristic of the chondrichthyan endoskeletons, the data presented here indicate that this type of mineralisation has been significantly modified within the holocephalans. The mineralised components of the endoskeleton of *Callorhinchus milii* (Family Callorhinchidae) consist of small units that form a layer of tightly arranged, irregularly shaped tesserae, also present in *Hydrolagus* (Family Chimaeridae). These tesserae in *Callorhinchus* and *Hydrolagus* differ in many respects from most shark and ray tesserae, being smaller and simpler, lacking features such as distinct cap and body zones, mineralised spokes between the tesserae and retention of lacunae housing vital chondrocytes. Nevertheless some similarities in development are present, such as the intra-chondrocyte septa that surround the chondrocytes early in the development of the tesserae, described above in *Callorhinchus* and the ray *Urobatis* (Dean et al. 2009; Seidel et al. 2016). Tesserae in sharks such as *Scyliorhinus* and *Notorynchus* may also lack some features seen in the other elasmobranchs (Debiais-Thibaud, 2019: fig. 6.3; Seidel et al. 2016: fig. 11A). Tesserae in stem group holocephalans, as well as in fossil relatives of *Callorhinchus* such as *Edaphodon*, within the Family Callorhinchidae (Fig. 9F), also appear to possess the polygonal shape more characteristic of ray tesserae with these being larger and better developed than the mineralisation in the adult of *Callorhinchus*. Thus it appears that these smaller units may be the characteristic mineralised structure in extant holocephalans, representing a reduction of mineralisation occurring separately within the Callorhinchidae and Chimaeridae, and within the Elasmobranchii.

**FIGURE 11.**
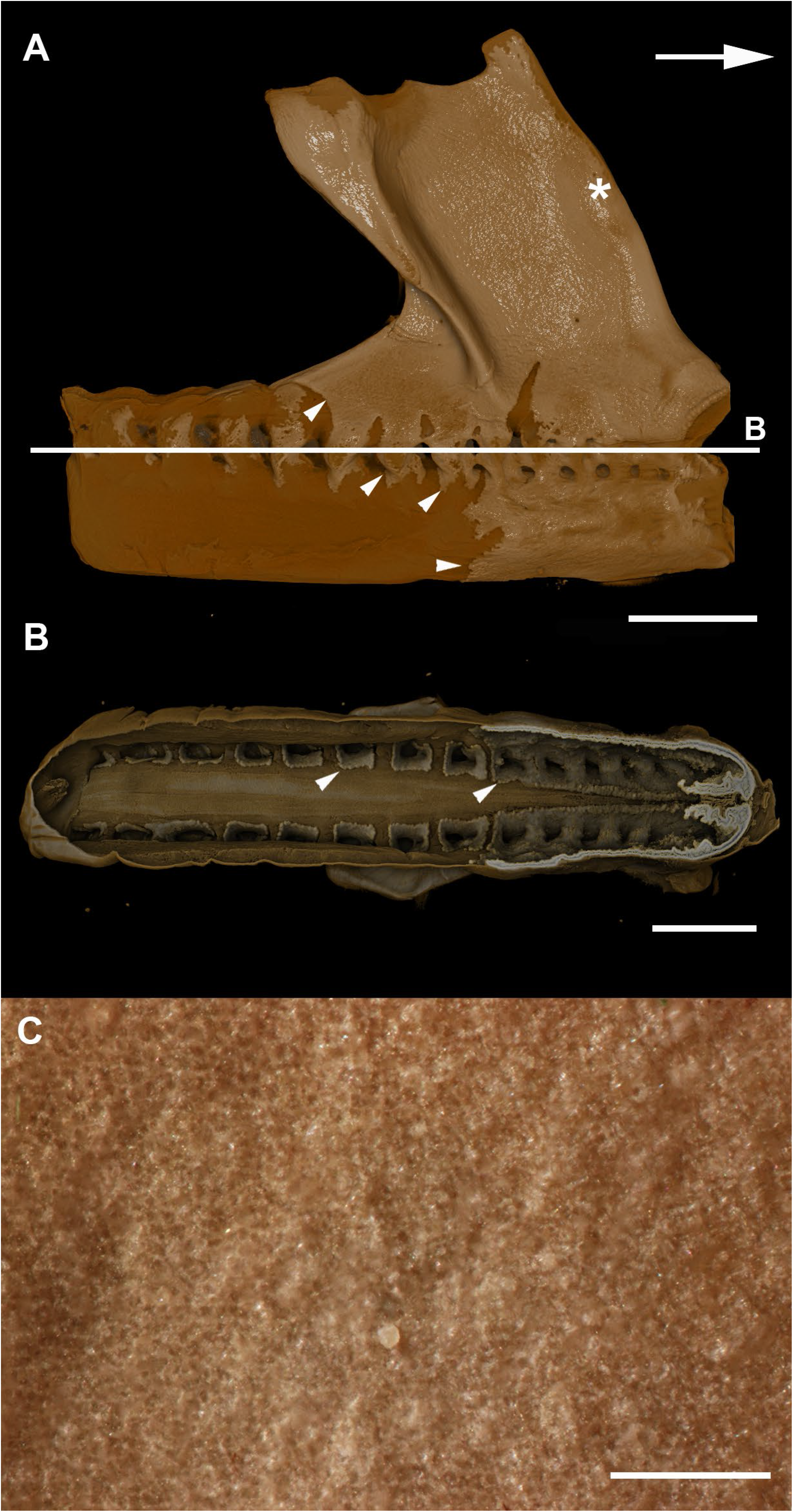
A, B, micro–CT scan of a synarcual from an adult *Callorhinchus milii* (Holocephali; Callorhinchidae). A,synarcual, lateral view; B, ocronal section (virtual) through synarcual; C, macrophotograph of lateral synarcual surface showing mineralisation with a granular appearance.

## 7 Author Contributions

ZJ, CB, JP conceived this project, JP, CB, ZJ contributed data to the project from fossil and extant holocephalans; all authors contributed to interpretation of the data and writing of the manuscript.

## 9 Acknowledgements

We would like to thank Ollie Crimmen and James MacLaine (NHM Life Sciences Department) for providing access to the slides of the *Callorhinchus* embryo and Innes Claxworthy (NHM Core Research Labs) for SEM imaging of the *Callorhinchus* adult. We would also like to acknowledge Elaine Miller and the John de Laeter Centre at Curtin University for expert advice, assistance, and service in the imaging of *Callorhinchus* embryo tissues, and the Curtin Faculty of Science and Engineering, School of Molecular and Life Sciences and Centre of Health and Innovation Research for support. The John de Laeter Centre is funded by the Australian Research Council (ARC LE130100053). JP is supported by the Australian Government through the AGRTP Scholarship. We would also like to thank Alan Pradel and John Maisey for discussion of tesserae in fossil chondrichthyans.

