## Supplemental Figure 1 for "Mineralisation of the *Callorhinchus* vertebral column (Holocephali; Chondrichthyes)"

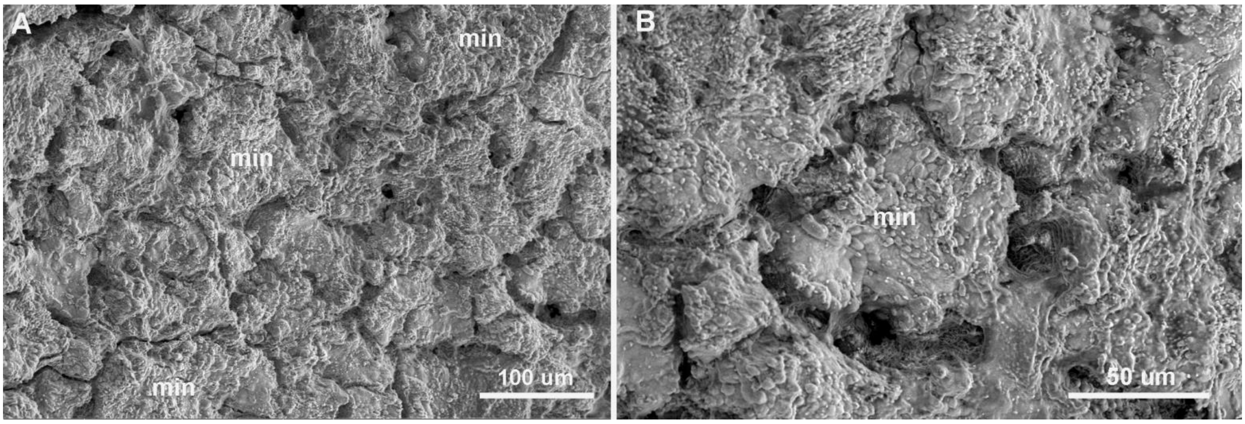


SUPPLEMENTARY INFO FIGURE 1. SEM images of mineralisation from the synarcual (anterior fused vertebrae) of the second adult *Callorhinchus milii* (Holocephali; Callorhinchidae). A, tesselated mineralisation from a planar perspective; B, close up of mineralisation from a planar perspective.
